# Molecular profiling in HNSCC patients targets potential responsiveness to ICIs

**DOI:** 10.1101/2022.12.16.520703

**Authors:** Andrea Sacconi, Paola Muti, Claudio Pulito, Raul Pellini, Sabrina Strano, Paolo Bossi, Giovanni Blandino

## Abstract

Immunotherapy is emerging as a valid therapeutic strategy for various cancers although only a subset of patients responding to the therapy. The response to immunotherapy remains low for many cancer types because of poor ability to appropriately classify responding patients.

We analyzed TCGA cohort of HNSCC patients in relation to a 26 immune gene set and cell types to define signaling pathways associated with resistance to immune checkpoint blockade. Results were validated on a cohort of 102 HNSCC patients under treatment with PD-L1 inhibitors and by *in vitro* experiments.

Using bioinformatic approach, we observed a significant association between the gene set and TP53 gene status and other predictors of ICIs’ response in HNSCC patients. The presence of both TP53 mutation and co-mutations was associated with significantly higher levels of the immune gene expression than in lesions with only the TP53 mutated. We also observed that higher level of a MYC signature was associated with lower level of the immune gene expression. *In vitro* and cohort validation corroborated the evidence.

**SIGNIFICANCE:** Immune gene signature sets represent biomarkers to be implemented for the better classification of those HNSCC patients responsive to immunotherapy. A particular advantage of these biomarkers is the ability to characterize HNSCC patients using tumor tissues easily applicable in clinical setting.

## Introduction

Every year, almost one millions of people are affected by Head and Neck Squamous Cell Carcinoma (HNSCC) in the world (1). HNSCC is composed by a biologically diverse and genomically heterogeneous disease that emerges from the squamous mucosal lining of the upper aerodigestive tract, including the lip and oral cavity, nasal cavity, paranasal sinuses, nasopharynx, oropharynx, larynx and hypopharynx (2). Most patients present with locally advanced disease with a high risk of recurrence, and approximately 10% of them presents with metastatic disease (3). The 5-year survival for HNSCC patients across all stages of 40%–50% for cancers determined with the median overall survival for recurrent/metastatic (R-M) patients of 10–13 months (4). In 2016, the US Food and Drug Administration (FDA) approved two immunotherapeutic agents, the anti-programmed cell death protein (PD-1) monoclonal antibodies, nivolumab (Opdivo, Bristol-Myers Squibb) and pembrolizumab (Keytruda, Merck), for the treatment of R-M HNSCC patients refractory to platinum-based therapy. The same agency approved then pembrolizumab for the first-line treatment of patients with unresectable R-M HNSCC. Immune checkpoint inhibitors (ICIs) are an active category of immunotherapies that block inhibitory immune check-point pathways in order to reactivate immune responses against cancer.

The use of immune checkpoint inhibitors (ICIs) is increasing in several cancer settings, both as monotherapy and as combinations with another ICI, chemotherapy, or targeted agents. The benefit of this new class of drugs seems to be large but limited to a subgroup of patients, thus an efficient patient characterization is needed to guide improvements in treatment. Predictors of response to ICIs are critical to ensure correct selection of patients to be offered immunotherapy, thus achieving higher response, preserving patients from unnecessary toxicities, and saving economical resources. However, till now there are limited markers available, mainly consisting in PDL1, microsatellite instability and tumour mutational burden (TMB) assessment. Several clinical, molecular, and microbiological factors are assumed to have a role as influencing response to ICIs.

The aim of the present report is to assess the role of TP53 gene status and other characteristics as predictors of ICIs’ response in HNSCC patients, adjusting for potential confounders and using a bio-informatic approach which results have been validated by ad-hoc conducted experimental studies and corroborated by analyses carried out in cohorts of HNSCC patient.

## Results

The 520 participants of the TCGA sample were equally distributed for age, gender (males/females) and smoking. 19% (N=97) of the patients were HPV positive and, as expected, only 4% (N=20) had a stage 1 lesion at diagnosis. Most patients had cancer lesions characterized by mutated TP53 and wild type CDKN2A, FAT1 and PIK3CA genes (data are reported in **Table A** only for reviewers’ perusal). The 26 immune gene set, the related analysis of the 150 gene expression together with the corresponding 15 immune cell types are described in **Table S1**.

To test the hypothesis that TP53 gene status and other characteristics could represent predictors of ICIs’ response in HNSCC patients, we used a step-by-step approach depicted the **Figure 1** flowchart. **The Step 1** of the flowchart includes mainly database analyses. Phase **A)** describes the analysis of 26 immune genes signature as prognosis predictor in the cohort of 520 HNSCC TCGA. **Phase B**) Analysis of the signature prediction performance by HNSCC TCGA subgroups such as TP53 wild type versus mutated and TP53 co-mutated. **Phase C**) Analysis of signature association with 22 MYC-related genes expression and cell type enrichment. In our previous work we demonstrated the role of a 22 genes MYC-related signature in HNSCC cancer as surrogate of TP53 mutation (5). In **Step 1**, also includes an in-vitro validation of Phase B and Phase C study results (Fig S3). In **Step 2**, we conducted a general result validation on 102 HNSCC patients of the GEO database.

**Fig 1:**
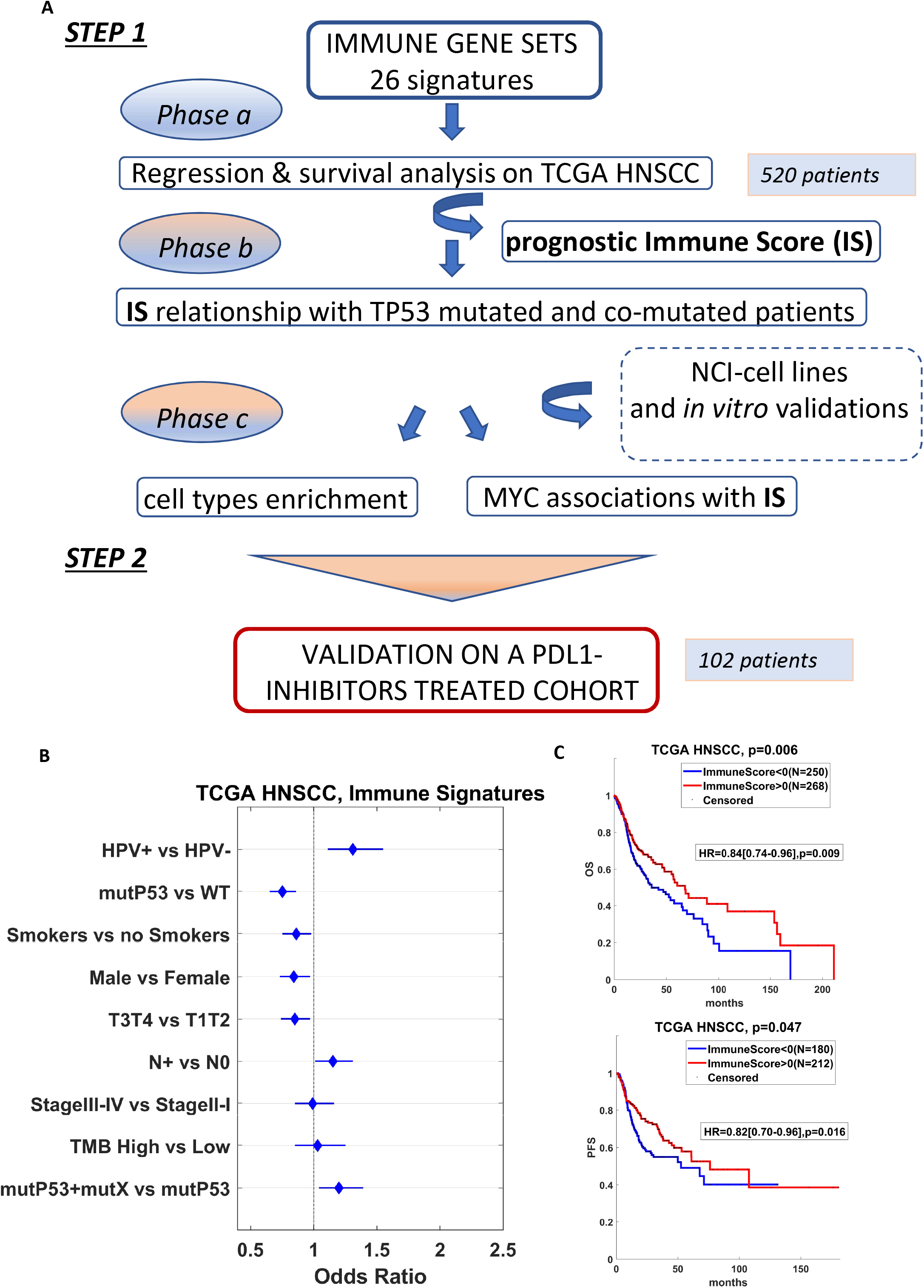
Regression and survival analysis on TCGA HNSCC cohort. **A)** Workflow of the analyses. **B)** Forest plot representing the association of average expression of 150 genes included in the 26 immune gene sets and the clinical variables in 520 HNSCC patients from TCGA. Results of the linear regressions are shown as Odds Ratio with confidence intervals at 95%. **C)** Kaplan-Meier curves of HNSCC patients from TCGA cohort with high or low Immune Scores evaluated for overall survival and progression free survival (upper and bottom panel, respectively). Differences between curves were evaluated by logrank test. Hazard ratios with 95% confidence intervals were assessed by Cox Hazard regression models. Immune Scores were evaluated as the positive and negative z-scores of the average expression of the 150 genes composing the immune gene sets.

### STEP 1 – Regression and survival analysis

In **Figure 1**, panel **B**, we present the first set of results represented by the forest plot with Odds ratio with 95% CI of demographic and prognostic predictors of the immune signature expression by using regression models in HNSCC dataset from TCGA. All the analyses were performed at univariate level considering the effect of only one independent variable (demographic or prognostic variables) on the signature (dependent variable). We observed that only the **HPV** status, with HPV negative versus the positive lesions, **lympho-node status** (N0 vs N+) and **TP53** mutated plus additional mutations versus only TP53 mutated lesions, were statistically associated with a higher signature expression. Details of regression analyses on clinical factors for each immune gene set are shown in Fig. S1 and Fig. S2. Notably, building a multivariable regression model, TP53 mutational status resulted the only clinical factor significantly associated to the immune signature (Table S2). To provide clinical meaning to the described results, in **Figure 1, panel C**, we reported overall-survival (OV) and progression-free-survival curves assessed in the TCGA cohort of 520 patients. In that cohort high immune signature expression was associated with a statistically significant longer survival.

Because the mutational TP53 status was so important, we assessed the expression distribution of the immune signature by three groups of patients with wild type TP53 status, TP53 mutant status as well as TP53 mutant plus one of the other three most frequent mutations observed in HNSCC cancer patients (FAT1, CDKN2A, and PI3K genes). Wild type TP53 patients were characterized by a higher signature expression level compared with those carrying TP53 mutation and TP53 co-mutations (TP53 mutation/FAT1, TP53/CDKN2A, TP53/PI3K) and the last group had an immune gene expression higher than the TP53 mutated one (Figure 2, panel A).

**Fig 2:**
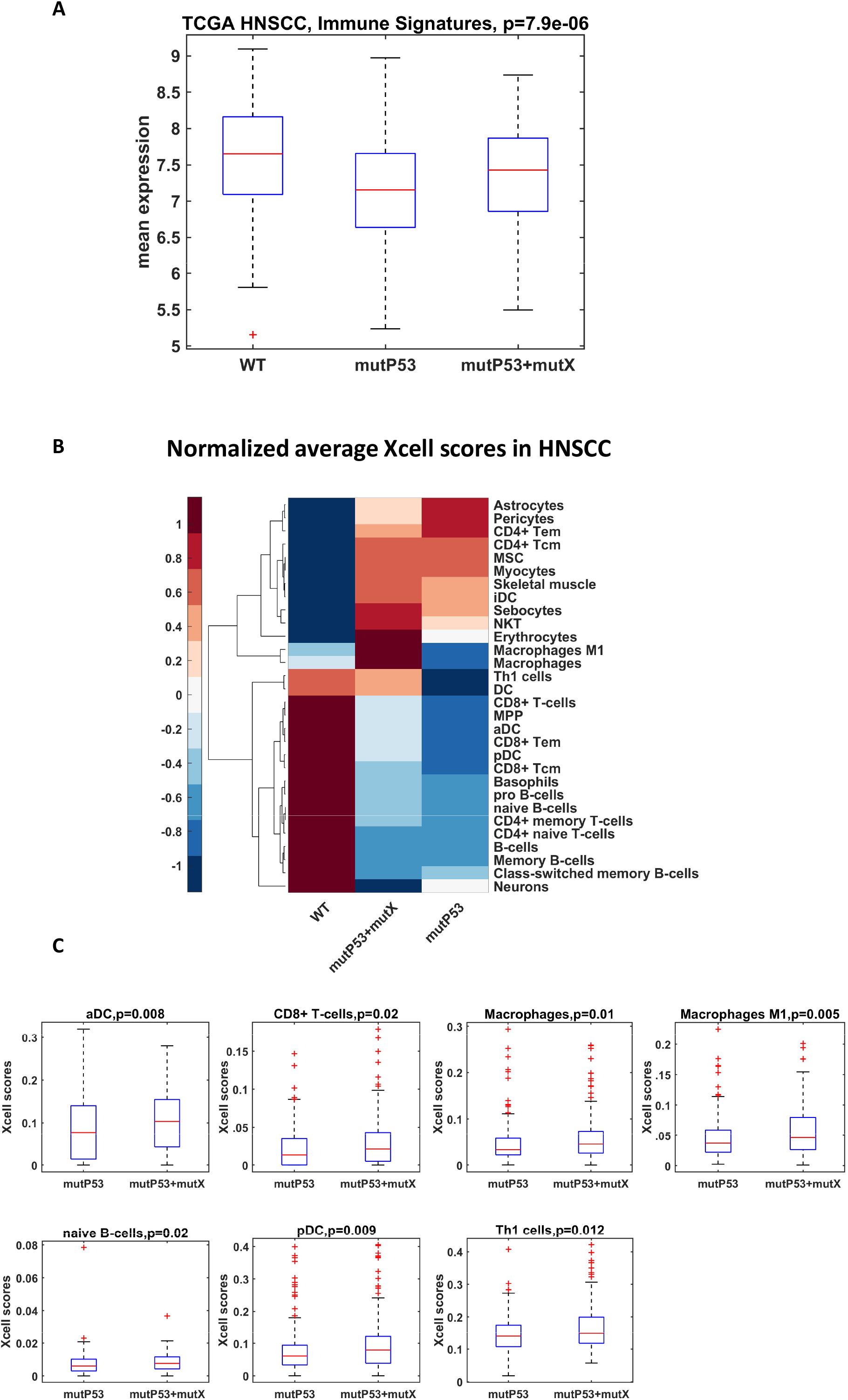
Evaluation of immune gene sets expression and cell type enrichments reveals differences between TP53 mutated patients and co-mutated patients. **A)** Distributions of the gene signature composed by the average expression of 150 genes of the immune gene sets by TP53 mutation and TP53 mutation carried on other mutations among FAT1, CDKN2A and PIK3CA in HNSCC patients (106 WT, 171 TP53 and 189 TP53+mutX). P-values were evaluated by KruskalWallis test. **B)** Cell types enrichment analysis by comparing 64 cell type signatures in subgroups of HNSCC patients with TP53 mutation, TP53 mutation with other mutations and wild type patients. Heatmap representing the normalized average scores obtained from Xcell software, reflecting the cell type abundance of the most significant modulated cell types among the three subgroups. The statistical significance (p<0.05) was assessed by KruskalWallis test. **C)** Cell types enrichment of TP53 mutated patients and TP53 mutated patients who harboured other mutations. Scores were obtained from Xcell software. P-values were evaluated by Wilcoxon ranksum test.

These signatures were validated using extensive in-silico simulations and cytometry immunophenotyping. **Panel B**, Figure 2, displays results of a deconvolution analysis with cellular heterogeneity landscape of tissue expression profiles by an adjusted enrichment score (XCell score). The scores represent a measure of cell type abundances, enabling comparison across cell types and across samples (6). The analysis identified the lineage marker genes by TP53 wild type, TP53 mutated and co-mutated patients. The 30 cell types reported on the side of the heatmap had relative cell types abundances differently distributed across the three group of TP53 status and TP53 co-mutated patients. Finally in **panel C**, we reported only the cell types specifically modulated between TP53 mutated and TP53 co-mutated.

To detail functional connection between gene mutation and immune signature, we assessed the role of the MYC-signature. In **Figure 3**, we assessed the expression of PDL1 and CTL4 in TCGA patients with high or low expression of MYC-related signature in TCGA (panel A and B) and the expression of the immune gene sets in both TCGA and GEO cohort (GSE195832) (panel C and D) patients. Again, lower expression level of the MYC-dependent signature was significantly associated with higher levels of immune gene sets, PDL1 and CTLA4. For validation purpose, we also performed qRT-PCR analysis of PD-L1 (Fig.3 panel E) and CTLA4 (Fig.3 panel F) in Cal27 treated with JQ-1. The latter is a small-molecule that inhibit the activity of the BET family proteins by masking their bromodomain acetyllysine-binding pockets (7). JQ-1 has been demonstrated to act as antineoplastic agent by mainly inhibiting c-MYC functions Both genes showed increased expression after treatment when compared to their controls, strengthening the potential role of MYC in an immunogenic context.

**Fig 3:**
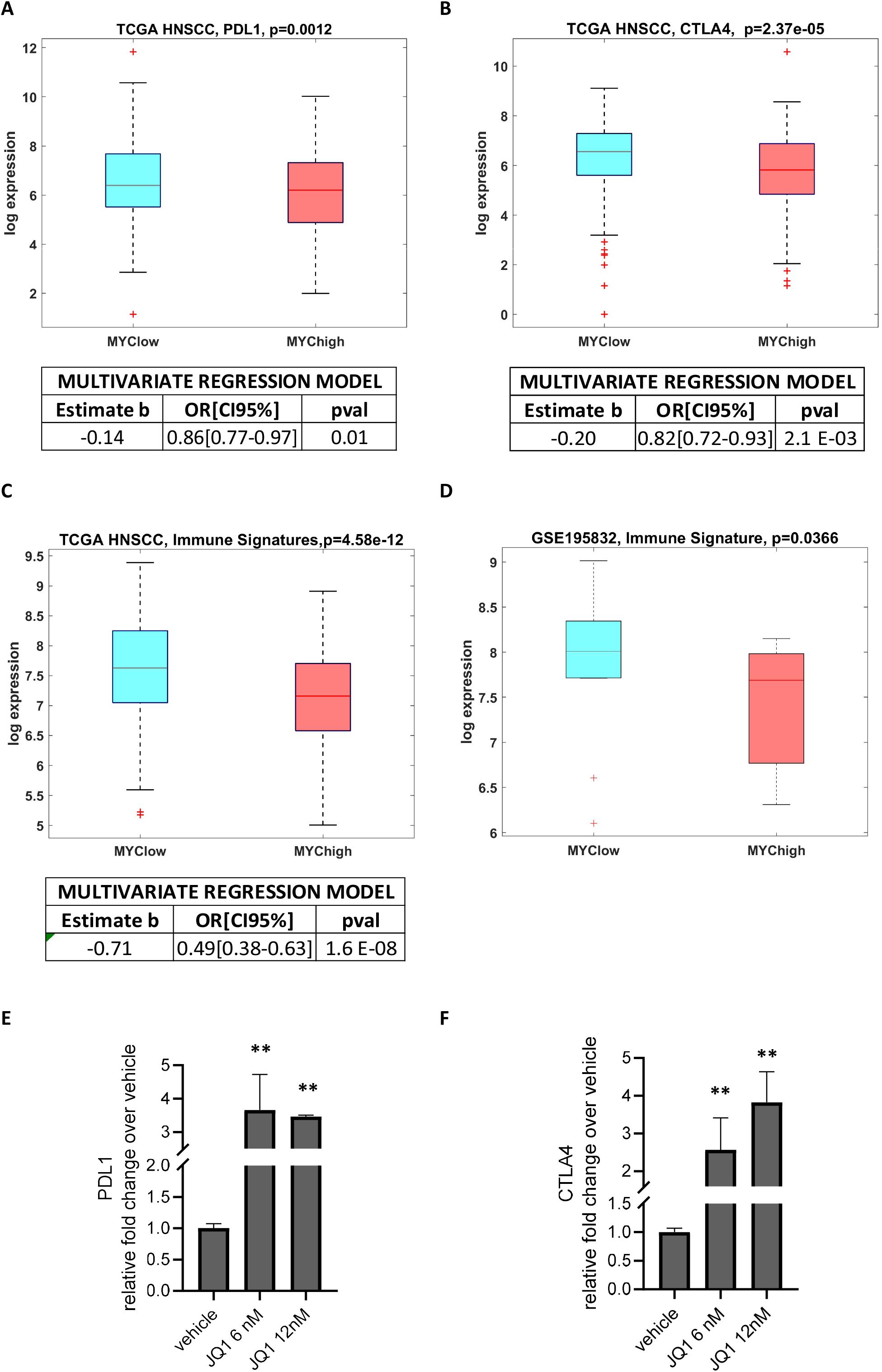
Association of a MYC dependent signature with the immune gene sets. **A-C)** box-plot of the PD-L1 (A), CTLA4 (B) expression values and mean expression of 26 immune gene sets (C) in patients with high and low level expression of a 22 genes signature MYC dependent in HNSCC datasets from TCGA. Statistical significance between distributions was assessed by Wilcoxon rank-sum test. Multivariate regression models were built to adjust the differences of the genes between patients with high and low MYC signature. The models include T status, TP53 mutation, gender, smoking status and, HPV status. High and low expression of the MYC signature were evaluated by positive and negative z-scores of the mean gene expression, respectively. **D)** box-plot of the mean expression of 26 immune gene sets in 28 pre-treated HNSCC patients with high and low level expression of a 22 genes signature MYC dependent in GSE195832 dataset from GEO. Statistical significance between distributions was assessed by Wilcoxon rank-sum test. **E-F)** qRT-PCR analysis of PD-L1 (E) and CTLA4 (F) in Cal27 treated with JQ-1. Bars indicate the average of at least three independent experiments. Statistics (t-test): * p< 0.01, ** p<0.005.

### Step2: Analysis of the immune gene sets and MYC dependent signature in a cohort of HNSCC patients treated with PDL1-inhibitors

In 2022, Foy and colleagues published data reporting analysis conducted in 102 and 82 patients with HNSCC or NSCLC in which they defined an immunologically active phenotype score by gene expression profiling (8). In that publication, the authors reported that a high 27-gene expression, different from our signature, defined as ‘HOT’ score was associated in treated with PD-1/PD-L1 inhibitors with a significant improved OS and PFS.

Thus, coherently with their approach, we further investigate the role of the immune gene sets and of the MYC dependent gene signature in their well characterized cohort of HNSCC patients under treatment with PD-L1 inhibitors obtained from GEO database (accession ID: GSE159067).

In Fig. 4, we report results of our analysis on the prognostic value of the tested our immune gene sets in both OV (left panel) and PFS (right panel). Our results corroborated the previous evidence (Fig. 1, panel C) of a better clinical performance of high versus low signature expression.

**Fig 4:**
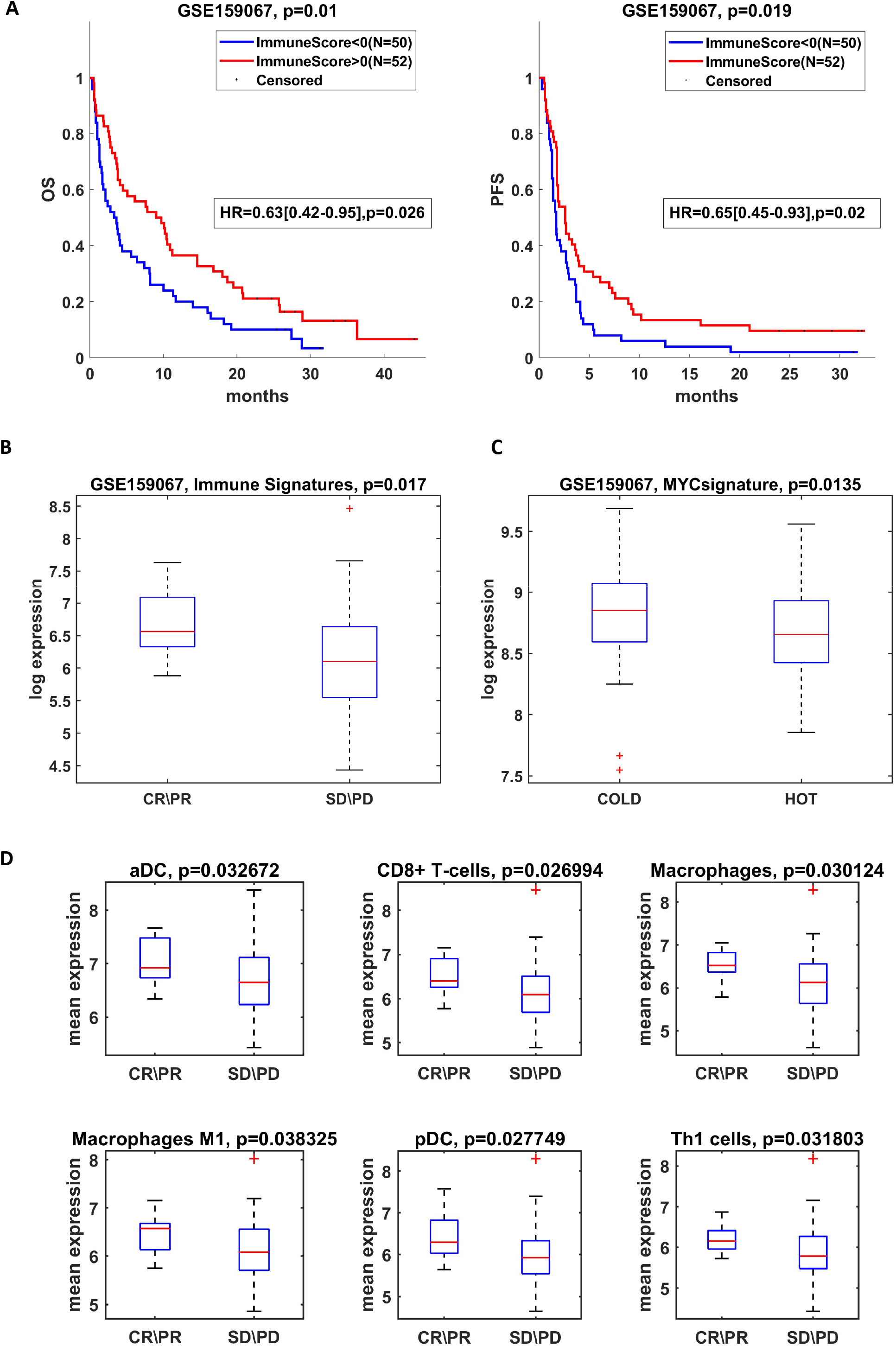
Validation of immune gene sets and MYC dependent signature in a cohort of HNSCC patients treated with PDL1-inhibitors. **A)** Overall survival (left panel) and Progression free survival (right panel) of 102 patients treated with PDL1 inhibitors from GEO database (GSE159067). Patients were split basing on the Immune Score. The high\low levels of Immune Score were obtained considering the positive and negative z-scores of the average expression of the 26 immune gene sets, respectively. Differences between curves were evaluated by logrank test. The multivariate Cox Hazard regression analysis was adjusted for gender and HPV status. **B-C)** Average expression of the 26 immune gene sets and MYC signature distribution in 102 patients treated with PDL1 inhibitors (GSE159067, panel B and C, respectively). The immune gene sets expression was evaluated in patients with complete or partial response and patients with stable disease or progression disease after treatment (B). The MYC signature expression was evaluated according to the phenotype classification (“COLD” and “HOT” patients) obtained from Foy JP and colleagues (C). Differences were evaluated by Wilcoxon test. D) Significantly modulated cell type marker genes in 102 patients from GEO database (GSE159067) among the 7 cell types previously identified in the deconvolution analysis of TCGA HNSCC data. Statistical significance between patients with complete or partial response and patients with stable or progression disease after treatment with anti-PDL1 was evaluated by Wilcoxon test.

The immune gene sets we used to define our immune score was also strongly correlated to the classification (“COLD” and “HOT”) introduced by Foy and colleagues (Fig S4, panel A). Notably, In Fig. 4, panel B, low levels of the immune gene sets are statistically significantly associated with stable or progression disease during immunotherapy. Furthermore, low level of the MYC dependent signature is resulted significantly associated with the immunologically “HOT” type (Fig. 4, panel C). These results are in line with our finding on TCGA data and cell lines about the potential role of MYC in an immunogenic context.

Finally, we used specific marker genes of the cell types identified in the cell enrichment analysis of TCGA data (Fig. 4, panel D) to evaluate their quantitative expression on immunotherapy treated patients. Six out of the seven investigated cell types resulted strongly up-regulated in HNSCC patients characterized by complete or partial response to the treatment. The same cell types showed a significant reduced abundance in patients with low Immune Score (Fig S4, panel B).

## Discussion

The successful implementation of precision medicine is highly based on clinically-relevant predictive biomarkers, a major challenge that is still being overcome. The administration of immune checkpoint inhibitors has significantly improved treatment outcomes and survival of HNSCC patients. However, a rather limited fraction of HNSCC patient benefits from ICIs treatment highlighting the unmet need to stratify better those patients to treat with immunotherapy (9). In the present work we applied a bio-informatic approach on a large-very-well characterized database such as the the Cancer Genome Atlas (TCGA) aiming to identify new biomarkers of immune modulation among the conventional risk factors or prognostic variables for patients affected with HNSCC. In HNSCC, other than PD-L1 for pembrolizumab, no additional biomarker has been identified until now. Also, the lack of immuno-modulatory effect of variables such as TMB, which appears important for the response to ICIs in other tumor types is a noteworthy aspect, as well as all the variables of tumor stage at diagnosis seem to lose relevance as biomarkers of potential response to ICI when looking at outcomes adjusted specifically for HPV and TP53 status. The application of 26 gene sets associated with immune functions to HNSCC mRNA expression data allowed identifying an immunoscore that distinguishes high versus low immunoscore patients. We also found that variables such HPV negative versus positive, and mutated TP53 versus WT-TP53 patients were associated with a higher immunoscore and a higher PDL-1 and CTLA4 expression. We also evidenced that HNSCC cell lines carrying WT-TP53 exhibit higher expression levels of both PDL1 and CTLA4 when compared to cell lines bearing TP53 mutations (S-Fig.3A). We have previously shown that gain of function activity of TP53 missense mutations in HNSCC also occurs through the aberrant transcriptional activation of a MYC-responsive 22 gene signature that is curtailed by the PI3K inhibitor alpelisib (5). We found that HNSCC patients with low expression of MYC signature expressed higher levels of both PDL1 and CTLA4 when compared with those expressing high levels of MYC signature. Interestingly, HNSCC patients with low expression of MYC signature have higher immunoscore. Depletion of either mutant p53 protein or its co-factor YAP released PDL1 expression in HNSCC cell lines. Of note, HNSCC patients carrying co-mutations such TP53/FAT1, TP53/CDKN2A, TP53/PI3K exhibited a higher immunoscore than those with only TP53 mutations. PDL1 expression levels were higher in HNSCC cell lines carrying TP53 co-mutations compared to those carrying only TP53 mutation (S-Fig.3B). The treatment with Alpelisib as a selective inhibitor of p110α-subunit of PI3K in CAL-27 and Detroit-562 head and neck cell lines enhanced the expression of PDL1. There are few important implications emerging from these findings. Firstly, while TP53 gain of function mutant p53 proteins might directly repress the expression of ICs such as PDL1, the presence of a co-mutation mitigates this effect through a yet unidentified compensatory mechanistic which renders this subset of HNSCC patients with higher immunoscore than mut-TP53 patients. Secondly, HNSCC patients carrying co-mutations TP53/PI3K could benefit from Alpelisib plus anti-PDL1 immunotherapy. Thirdly, HNSCC patients relapsing to precision therapeutic targeting herein exemplified by the PI3K inhibitor alpelisib might be proposed for immunotherapy treatment. With the due limitations deconvolution analyses from bulk-RNA-Seq data revealed that high/low immunoscore might contribute to decipher the immune infiltration cellular landscape of HNSCC patients. Indeed, we found that high-immunoscore HNSCC patients exhibited immune infiltration in which aDC, macrophages, CD8+T-, macrophages M1, naïve B cells, pDC, Th1 cells appear to be significantly more represented than in those with low immunoscore. Notably, HNSCC patients with TP53 and additional mutations showed a putative immuno cellular landscape more similar to WT-P53 HNSCC patients than to those with TP53 mutations. There are no current therapies that target directly infiltration immune cells for HNSCC patients, but the potential of the identified immunoscore to provide insights on the cellular composition of the immune infiltrate is certainly relevant to profile immunologically a given patient. So far, the identified immunoscore holds prognostic value as those HNSCC patients with low immunoscore exhibited shorter OS and PFS than high-score patients and allows identifying deeply HNSCC patients who could potentially benefit from immunotherapy. We subsequently tested the predictive potential of the identified immunoscore using expression data of a cohort of 102 HNSCC patients treated with PDL1 inhibitors (8). The classification of HNSCC treated patients following high versus low immunoscore revealed that those with low immunoscore have shorter OS and PFS than HNSCC patients with high immunoscore. In aggregate, our findings provide evidence of an immunoscore which holds prognostic and predictive features of a biomarker to select HNSCC patients to be treated with immunotherapy.

## Methods

The study was conducted to investigate the association between 26 immune gene sets (listed in **Table S1**) and immune check point proteins expressed in HNSCC and their effect on patient survival. The 26 immune gene-set signature tested in the present report was identified in a previous publication where it was found associated with ICIs’ response in Triple Negative Breast Cancer (10). In this study, we applied the signature on HNSCC as the 20 genes of the 26 included in the signature were also found modulated in HNSCC (11). In particular, in the present analysis we assessed the effect of the immuno-signature on the two most known immune checkpoint proteins validated for clinical use also in HNSCC: the programmed cell death protein ligand - 1 (PDL-1) and the T-lymphocyte associated protein 4 (CTL-4) controlling for potential confounders and effect modifiers (12). Data derived from the “The Cancer Genome Atlas” (TCGA - the HNSCC-TCGA, Nature 2015) and the analyses included 520 HNSCC patients. We gathered the normalized TCGA HNSCC gene expression of tumour from Broad Institute TCGA Genome Data Analysis Center (http://gdac.broadinstitute.org/): Firehose stddata 2016_01_28 and Broad Institute of MIT and Harvard. <u>doi:10.7908/C11G0KM9.</u> Clinical information for cohorts was collected from cBioPortal (https://www.cbioportal.org/datasets) according to the data published by Liu et al. (10).

To identify HNSCC patients responsive to immunotherapy, for each gene set and immune checkpoint protein we initially developed logistic regression model based on the average expression of the immune signature genes. Specifically, the mean expression values of the genes belonging to specific immunological signature were then used to build linear regression models and to assess their associations with several clinical variables. Odds ratio with confidence intervals at 95% were evaluated for each gene set by including age, gender, T, N, stage, HPV status, smoking history, TMB, and the mutational status of TP53, **PIK3CA, FAT1, CDKN2A**. Significance was defined at the 5% level. Results are represented by Odds Ratio values (OR) with the 95% intervals of confidence. Significance of genes modulation between different subgroups of samples was assessed by Wilcoxon test and ANOVA test. The analyses were conducted with Matlab R2020b.

Kaplan-Meier curves of HNSCC patients with high or low immune scores were conducted to assess the overall survival (OS) and progression free survival (PFS) differences between curves were evaluated by logrank test. Hazard ratios with 95% confidence intervals were assessed by Cox Hazard regression models. Immune scores were evaluated as the positive and negative z-scores of the average expression of the 150 genes composing the immune gene sets.

To investigate the cellular heterogeneity landscape of the tissue expression profiles, we performed a cell type enrichment analysis by using *XCell* (https://xcell.ucsf.edu/). The tool is a gene-signature based method able to reach reliable associations in the gene expression of 64 immune and stroma cell types (6).

A validation cohort of 102 HNSCC patients treated with PDL1 inhibitors was gathered from GEO database with accession ID GSE159067 (8).

## Authors’ Contributions

Conception and design: A.S., P.B., G.B. Data analysis A.S. Interpretation: A.S., P.M., S.S., P.B., G.B. In vitro experiment: C.P. Manuscript writing and revision: A.S., P.M., R.P, S.S., P.B., G.B. Final approval of manuscript: All authors.

## Figure legend

**Table A:** descriptive characteristics of TCGA HNSCC dataset.

**Table S1:** list of the immune gene sets analysed.

**Table S2:** Multivariate regression models of the main clinical variables associated with the immune signatures in HNSCC. The model was built considering a gene signature including all the 150 genes composing the immune gene sets.

**Supplementary Fig S1: A-D)** Forest plot representing Odds ratio with 95% CI of clinical predictors of several immune cell types and functional gene sets by using regression models in HNSCC dataset from TCGA. Red line highlights the behaviour of PD-L1 for comparison with other gene sets. All lines that don’t cross the 1 value are statistically significant. Each variable was dichotomized in the models to compare subgroup of patients by HPV status (A), tumor mutational burden (TMB) (B), and TP53 mutational status in concomitance or not with other mutations among FAT1, CDKN2A, PIK3CA (mutX) (b and c, respectively).

**Supplementary Fig. S2: A-F**) Forest plot representing Odds ratio with 95% CI of clinical predictors of 26 immune cell types and functional gene sets by using regression models in HNSCC dataset from TCGA. Red line highlights the behaviour of PD-L1 for comparison with other gene sets. All lines that don’t cross the 1 value are statistically significant. Each variable was dichotomized in the models to compare subgroup of patients by gender (A), smoking history (B), tumor size (C), lympho-node status (D) and stage (E).

**Supplementary Fig. S3: A)** Heatmap of PDL1 and CTLA4 expression from 15 HNSCC cell lines harbouring TP53 mutation and 3 WT cell lines obtained from NCI-60 cell lines dataset (left panel), and the relative box-plots of the distributions (right panel). Differences were evaluated by Wilcoxon test. **B-D)** qRT-PCR analysis of PD-L1 in Cal27, FaDu and Detroit 562 cell lines (b), in Cal27 depleted or not of p53 and YAP (c), or treated with 5nM of Byl-719 (d). Statistics (t-test): * p< 0.01, ** p<0.005

**Supplementary Fig. S4: A)** Average immune gene sets distribution in 102 patients treated with PDL1 inhibitors (GSE159067) according to the phenotype classification (“COLD” and “HOT” patients) obtained from Foy JP and colleagues (Ref). Differences were evaluated by Wilcoxon test. B) Significantly modulated cell type marker genes in 102 patients from GEO database (GSE159067) among the 7 cell types previously identified in the deconvolution analysis of TCGA HNSCC data. Statistical significance between patients with high and low Immune Score was defined as z-score of the average expression of the 26 immune gene sets.

