## Supplementary Figures for "Molecular profiling in HNSCC patients targets potential responsiveness to ICIs"

**Figure S1**

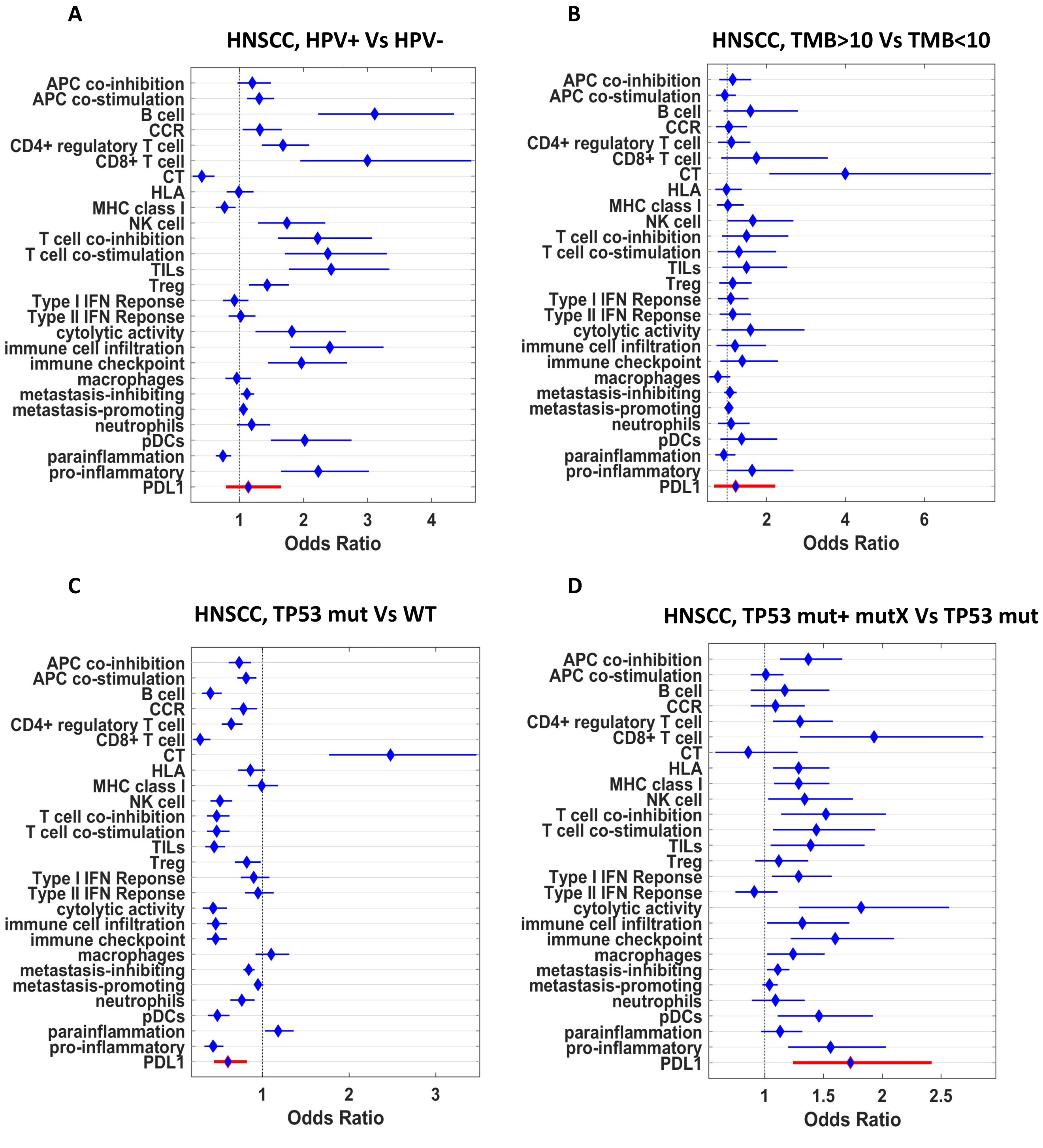
A
HNSCC, HPV+ Vs HPV-
B
HNSCC, TMB>10 Vs TMB<10
C
HNSCC, TP53 mut Vs WT
D
HNSCC, TP53 mut+ mutX Vs TP53 mut
APC co-inhibition
APC co-stimulation
B cell
CCR
CD4+ regulatory T cell
CD8+ T cell
CT
HLA
MHC class I
NK cell
T cell co-inhibition
T cell co-stimulation
TILs
Treg
Type I IFN Reponse
Type II IFN Reponse
cytolytic activity
immune cell infiltration
immune checkpoint
macrophages
metastasis-inhibiting
metastasis-promoting
neutrophils
pDCs
parainflammation
pro-inflammatory
PDL1
Odds Ratio

Figure S2

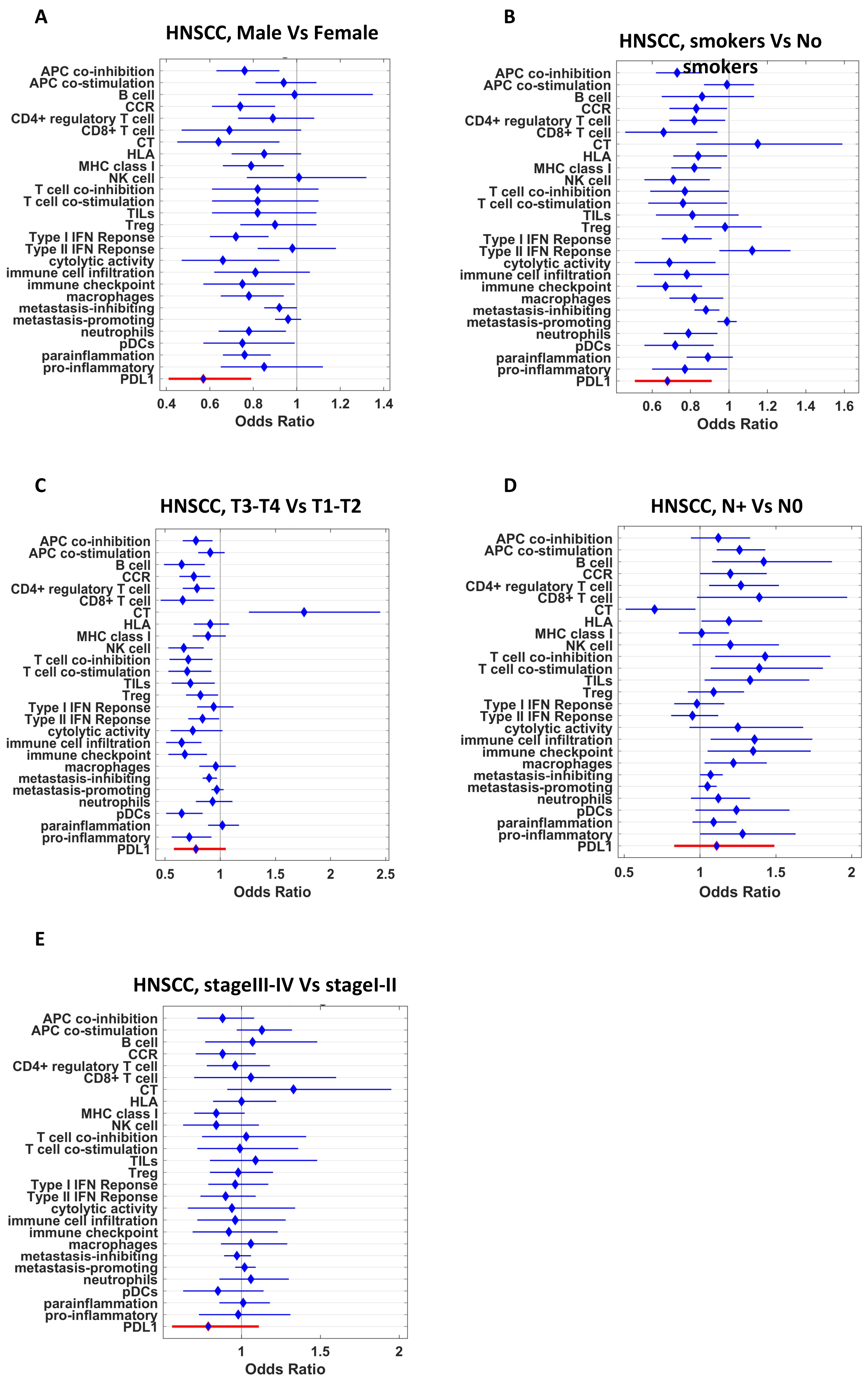
A
HNSCC, Male Vs Female
APC co-inhibition
APC co-stimulation
B cell
CCR
CD4+ regulatory T cell
CD8+ T cell
CT
HLA
MHC class I
NK cell
T cell co-inhibition
T cell co-stimulation
TILs
Treg
Type I IFN Reponse
Type II IFN Reponse
cytolytic activity
immune cell infiltration
immune checkpoint
macrophages
metastasis-inhibiting
metastasis-promoting
neutrophils
pDCs
parainflammation
pro-inflammatory
PDL1
0.4
0.6
0.8
1
1.2
1.4
Odds Ratio
B
HNSCC, smokers Vs No smokers
APC co-inhibition
APC co-stimulation
B cell
CCR
CD4+ regulatory T cell
CD8+ T cell
CT
HLA
MHC class I
NK cell
T cell co-inhibition
T cell co-stimulation
TILs
Treg
Type I IFN Reponse
Type II IFN Reponse
cytolytic activity
immune cell infiltration
immune checkpoint
macrophages
metastasis-inhibiting
metastasis-promoting
neutrophils
pDCs
parainflammation
pro-inflammatory
PDL1
0.6
0.8
1
1.2
1.4
1.6
Odds Ratio
C
HNSCC, T3-T4 Vs T1-T2
APC co-inhibition
APC co-stimulation
B cell
CCR
CD4+ regulatory T cell
CD8+ T cell
CT
HLA
MHC class I
NK cell
T cell co-inhibition
T cell co-stimulation
TILs
Treg
Type I IFN Reponse
Type II IFN Reponse
cytolytic activity
immune cell infiltration
immune checkpoint
macrophages
metastasis-inhibiting
metastasis-promoting
neutrophils
pDCs
parainflammation
pro-inflammatory
PDL1
0.5
1
1.5
2
2.5
Odds Ratio
D
HNSCC, N+ Vs N0
APC co-inhibition
APC co-stimulation
B cell
CCR
CD4+ regulatory T cell
CD8+ T cell
CT
HLA
MHC class I
NK cell
T cell co-inhibition
T cell co-stimulation
TILs
Treg
Type I IFN Reponse
Type II IFN Reponse
cytolytic activity
immune cell infiltration
immune checkpoint
macrophages
metastasis-inhibiting
metastasis-promoting
neutrophils
pDCs
parainflammation
pro-inflammatory
PDL1
0.5
1
1.5
2
Odds Ratio
E
HNSCC, stageIII-IV Vs stageI-II
APC co-inhibition
APC co-stimulation
B cell
CCR
CD4+ regulatory T cell
CD8+ T cell
CT
HLA
MHC class I
NK cell
T cell co-inhibition
T cell co-stimulation
TILs
Treg
Type I IFN Reponse
Type II IFN Reponse
cytolytic activity
immune cell infiltration
immune checkpoint
macrophages
metastasis-inhibiting
metastasis-promoting
neutrophils
pDCs
parainflammation
pro-inflammatory
PDL1
1
1.5
2
Odds Ratio

Figure S3

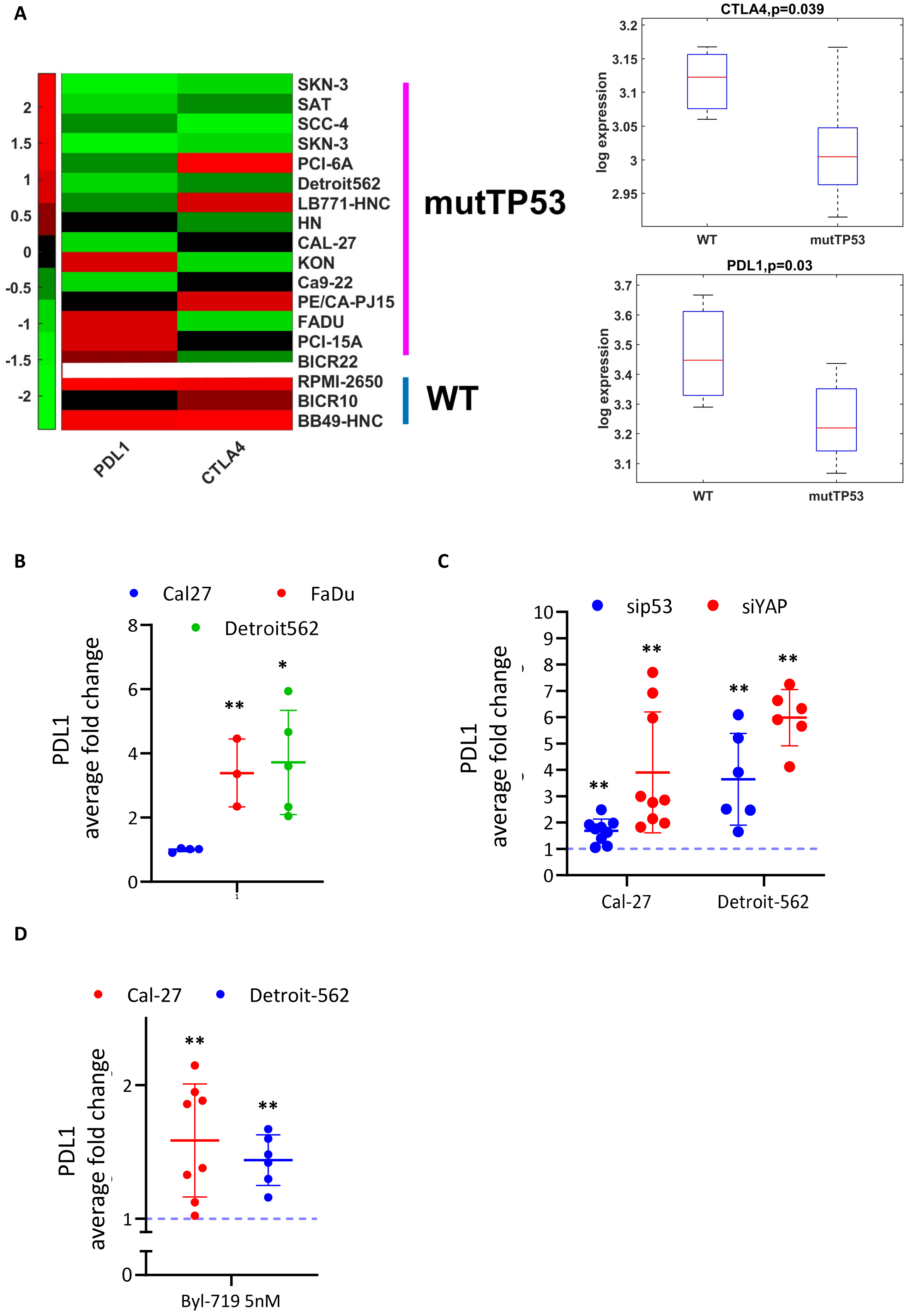
A
SKN-3
SAT
SCC-4
SKN-3
PCI-6A
Detroit562
LB771-HNC
HN
CAL-27
KON
Ca9-22
PE/CA-PJ15
FADU
PCI-15A
BICR22
RPMI-2650
BICR10
BB49-HNC
mutTP53
WT
PDL1
CTLA4
CTLA4,p=0.039
log expression
WT
mutTP53
PDL1,p=0.03
log expression
WT
mutTP53
B
Cal27
FaDu
Detroit562
PDL1
average fold change
C
sip53
siYAP
PDL1
average fold change
Cal-27
Detroit-562
D
Cal-27
Detroit-562
PDL1
average fold change
Byl-719 5nM

**Figure S4**

**A**

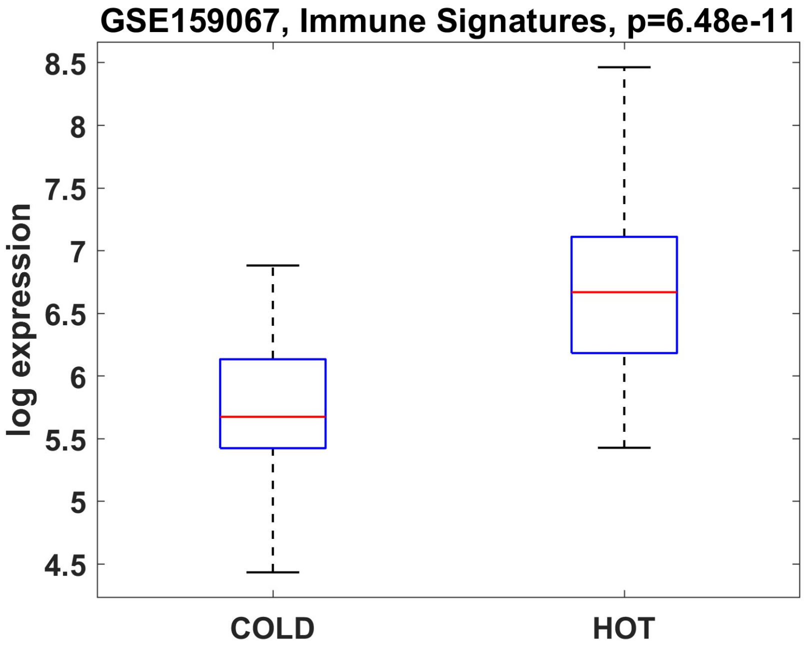
GSE159067, Immune Signatures, p=6.48e-11
log expression
8.5
8
7.5
7
6.5
6
5.5
5
4.5
COLD
HOT

**B**

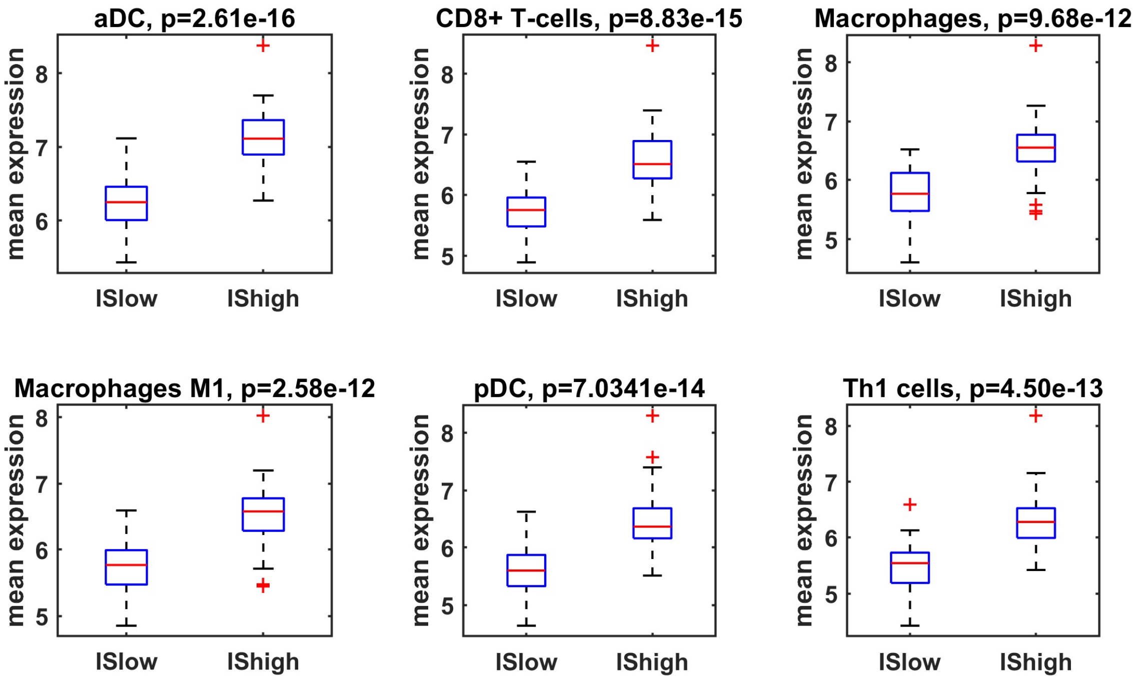
aDC, p=2.61e-16
CD8+ T-cells, p=8.83e-15
Macrophages, p=9.68e-12
Macrophages M1, p=2.58e-12
pDC, p=7.0341e-14
Th1 cells, p=4.50e-13
mean expression
ISlow
IShigh

**Table A**

| Variable | | #pz(%) |
|---|---|---|
| **Age** | age<=53(Q1) | 139(27%) |
| | 53<age<69(Q1-Q3) | 250(48%) |
| | age>=69(Q3) | 130(25%) |
| **Sex** | Female | 136(26%) |
| | Male | 384(74%) |
| **T** | T1 | 35(7%) |
| | T2 | 151(29%) |
| | T3 | 135(26%) |
| | T4 | 183(35%) |
| **N** | N0 | 244(47%) |
| | N+ | 258(50%) |
| **Stage** | StageI | 20(4%) |
| | StageII | 98(19%) |
| | StageIII | 105(20%) |
| | StageIV | 276(53%) |
| **HPV** | Neg | 421(81%) |
| | Pos | 97(19%) |
| **Smoking history** | No smokers | 117(22%) |
| | Smokers | 177(34%) |
| | Reformed smokers (>15years) | 73(14%) |
| | Reformed smokers (<15years) | 139(27%) |
| **Alchol consumption** | No | 162(31%) |
| | Yes | 347(67%) |
| **TP53 mutation** | WT P53 | 152(30%) |
| | MUT P53 | 360(70%) |
| **CDKN2A mutation** | WT CDKN2A | 400(78%) |
| | MUT CDKN2A | 112(22%) |
| **FAT1 mutation** | WT FAT1 | 399(78%) |
| | MUT FAT1 | 113(22%) |
| **PIK3CA mutation** | WT PIK3CA | 419(82%) |
| | MUT PIK3CA | 93(18%) |

**Table S1**

| | signature | genes |
|---|---|---|
| **15 immune cell types and function** | B cell | CD79B,BTLA,FCRL3,BANK1 |
| | CD4+ regulatory T cell | C15orf53,IL32,CTLA4,FOXP3 |
| | CD8+ T cell | CD8A |
| | NK cell | KLRF1,KLRC1 |
| | cytolytic activity | GZMA,PRF1 |
| | macrophages | CD68,CYBB,MMP9,LGMN |
| | MHC class I | HLA-A,B2M,TAP1 |
| | APC co-stimulation | ICOSLG,CD70,CD40,CD58 |
| | T cell co-stimulation | CD27,CD28,ICOS,CD2,CD226 |
| | APC co-inhibition | CD274,PDCD1LG2,C10orf54,LGALS9 |
| | T cell co-inhibition | CTLA4,LAG3,TIGIT,BTLA |
| | neutrophils | SELL,VNN3,KDM6B,MNDA |
| | pDCs | IRF8,GZMB,CXCR3,CLEC4C |
| | Type I IFN Reponse | MX1,MX2,ISG20,DDX4 |
| | Type II IFN Reponse | GPR146,SELP,AHR |
| | HLA | HLA-A,HLA-B,HLA-C,HLA-E,HLA-F,HLA-G,HLA-H,HLA-J,HLA-L,HLA-DMA,HLA-DMB,HLA-DOA,HLA-DOB |
| | CT | MAGEA2,MAGEB2,MAGEC2,PAGE4,PRAME,CTAG1B |
| | immune cell infiltration | FOXP3,CXCR5,CD68,CD247,CD8A,PTPRC,MS4A1,CD1A |
| | Treg | BCL11B,CD4,CCR8,FOXP3,MSLN,IL2RA,VTCN1 |
| | immune checkpoint | CTLA4,LAG3,IDO1,IDO2,TIGIT,BTLA,CD274,PDCD1LG2,PDCD1 |
| | TILs | CD2,CD6,CD8A,CD79A,CD247,CYBB,SELL,STAT4 |
| | CCR | CCR1,CCR2,CCR3,CCR5,CCR7,CCR8,CCR9,CSF2 |
| | metastasis-promoting | SPNS2,GRSF1,C17orf62,CYBB,FAM175B,BACH2,NCF2,ARHGEF1,FBXO7,TBC1D22A,ENTPD1,LRIG1,CYBA,HSP90AA1,NBEAL2 |
| | metastasis-inhibiting | IRF1,RNF10,PIK3CG,ATPBD4,SLC9A3R2,IRF7,FAM108A1 |
| | pro-inflammatory | STAT1,GZMB,CD19,CD8B,GNLY,IFNG,IL12A,PRF1 |
| | parainflammation | AIM2,CD14,CD276,HMOX1,LGMN,MX2,MMP7,TLR2 |

**Table S2**

| | HNSCC, average Immune signatures | | | | | |
|---|---|---|---|---|---|---|
| | Estimate b | OR[CI95%] | pval | Rsquare | adjRsquare | model pval |
| | | | | 0.055 | 0.047 | 1.1E-05 |
| **HPV+ vs HPV-** | 0.14 | 1.15[0.95-1.38] | 0.15 | | | |
| **mutP53 vs WT** | -0.23 | 0.79[0.67-0.92] | 0.003 | | | |
| **Smokers vs no Smokers** | -0.08 | 0.92[0.81-1.06] | 0.25 | | | |
| **Male vs Female** | -0.19 | 0.83[0.71-0.96] | 0.016 | | | |
| **T3T4 vs T2T1** | -0.09 | 0.91[0.80-1.05] | 0.19 | | | |
